# Super-enhancer switching drives a burst in germline gene expression at the mitosis-to-meiosis transition

**DOI:** 10.1101/2020.03.11.987180

**Authors:** So Maezawa, Masashi Yukawa, Xiaoting Chen, Akihiko Sakashita, Kris G. Alavattam, Matthew T. Weirauch, Artem Barski, Satoshi H. Namekawa

## Abstract

The testis has the most diverse and complex transcriptome of all organs due to bursts in expression of thousands of germline-specific genes. Much of this unique gene expression takes place when mitotic germ cells differentiate and enter into meiotic prophase. Here, we demonstrate that the genome-wide reorganization of super-enhancers (SEs) drives bursts of germline genes after the mitosis-to-meiosis transition. At the mitosis-to-meiosis transition, mitotic SEs dissolve while meiotic SEs are established. Meiotic SEs are associated with the activation of key germline genes, defining the cellular identity of germ cells. This SE switching is regulated by the establishment of meiotic SEs via A-MYB (MYBL1), a key transcription factor for germline genes, and by the resolution of mitotic SEs via SCML2, a germline-specific Polycomb protein required for spermatogenesis-specific gene expression. Prior to the entry into meiosis, meiotic SEs are preprogrammed in mitotic spermatogonia, serving to direct the unidirectional differentiation of spermatogenesis. We identify key regulatory factors for both mitotic and meiotic enhancers, revealing a molecular logic for the concurrent activation of mitotic enhancers and suppression of meiotic enhancers in the somatic and/or mitotically proliferating phase.

## Introduction

Meiosis is an essential step in the preparation of haploid gametes, and the transition from mitotic proliferation to meiosis is a fundamental event in the maturation of germ cells. In the mammalian male germline, this mitosis-to-meiosis transition coincides with a fundamental change to the transcriptome: a dynamic and massive change in genome-wide gene expression^1–4^. Due to the burst in expression of thousands of germline genes at the mitosis-to-meiosis transition, the testis has the most diverse and complex transcriptome of all organs^2,5,6^. Further, during spermatogenesis, progressive, dynamic chromatin remodeling takes place to produce haploid spermatids^7^. Together with genome-wide changes in gene expression, the mitosis-to-meiosis transition accompanies the dynamic reorganization of epigenetic modifications, accessible chromatin, and 3D chromatin conformation to prepare for the next generation of life^4,8–12^. Yet despite these recent advances in our understanding of the mitosis-to-meiosis transition, it remains largely unknown how DNA regulatory elements underlie the massive, dynamic transcriptional change in the male germline.

With this in mind, one class of DNA regulatory elements is of particular interest: enhancers. Enhancers play key roles in the control of cell type-specific gene expression programs through the binding of transcription factors (TFs) and interaction with promoters^13–15^. Mammalian cells contain thousands of active enhancers^16^. A portion of these enhancers cluster in aggregate to regulate the expression of genes important for establishing cellular identity^17,18^; these enhancer clusters have been termed ‘super-enhancers’ (SEs)^17^. SEs are prevalent in various cell and tissue types, and are also found in cancer cells, where they direct the expression of key tumor pathogenesis genes^19^. However, for the most part, the characterization of SEs has been limited to somatic and/or mitotically proliferating cells. Given the massive scale and scope of the mitosis-to-meiosis transcriptome change, there are compelling questions as to the existence of a uniquely meiotic type of SEs and, if present, to the function of SEs in the meiotic phase. Although a previous study suggested that enhancer activation is not involved in mouse spermatogenesis^8^, the detailed profiles of active enhancers remain undetermined in spermatogenesis.

Here, we determine the profiles of active enhancers in representative stages of spermatogenesis in mice and identify a meiotic type of SEs. We demonstrate that the switch from mitotic to meiotic types of SEs drives a dynamic change in the transcriptome at the mitosis-to-meiosis transition. This SE switching is regulated by the establishment of meiotic SEs via A-MYB (MYBL1), a key transcription factor for germline genes^20,21^, and by the resolution of mitotic SEs via SCML2, a germline-specific Polycomb protein required for spermatogenesis-specific gene expression^3^. We found that meiotic SEs are preprogrammed in undifferentiated spermatogonia prior to the mitosis-to-meiosis transition, suggesting that gene activation in meiosis takes place based on epigenetic mechanisms of preprograming. Through systematic analyses, we identified key regulatory factors for both mitotic and meiotic enhancers, thereby exposing the molecular logic of concurrent activation mechanisms for mitotic enhancers and suppression mechanisms for meiotic enhancers in the somatic and/or mitotically proliferating phase.

## Results

### The landscape of active enhancers during spermatogenesis

To determine the landscape of active enhancers in spermatogenesis, we performed chromatin immunoprecipitation with sequencing (ChIP-seq) for the histone modification H3K27ac, a marker of active enhancers^22^. We analyzed four representative stages of wild-type spermatogenesis: THY1^+^ undifferentiated spermatogonia, a population that contains both spermatogonial stem cells and progenitor cells; KIT^+^ differentiating spermatogonia from P7 testes; purified pachytene spermatocytes (PS) in meiotic prophase; and postmeiotic round spermatids (RS) from adult testes (Fig. 1a). We carried out H3K27ac ChIP-seq for two independent biological replicates, and we confirmed that ChIP-seq signals are consistent at genomic loci between the replicates (Fig. 1b, Supplementary Fig. 1). (While generated for and analyzed in this study, our H3K27ac ChIP-seq data for wild-type PS and RS was initially introduced in another study that analyzed active enhancers on the sex chromosomes^23^.) Consistent with the massive, dynamic transcriptional change occurring at the mitosis-to-meiosis transition, we observed H3K27ac peaks in mitotically proliferating spermatogonia (blue shadow), H3K27ac peaks unique to meiotic spermatocytes (red shadow), and constitutive peaks (gray shadow: Fig. 1b).

**Figure 1.**
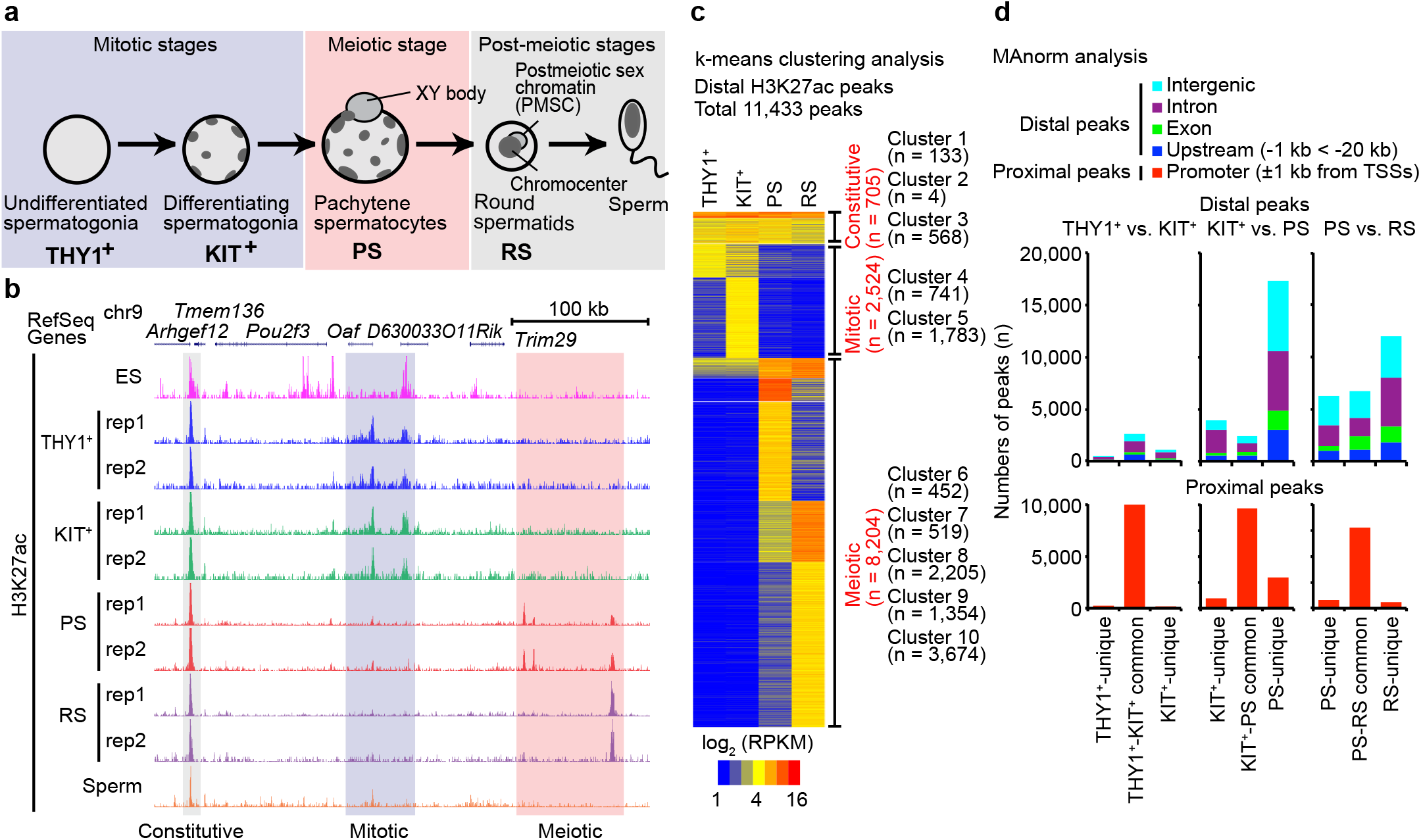
The landscape of active enhancers during spermatogenesis. (**a**) Schematic of mouse spermatogenesis and the five representative stages analyzed in this study: THY1^+^, undifferentiated spermatogonia; KIT^+^, differentiating spermatogonia; PS, pachytene spermatocyte; RS, round spermatids; Sperm, epididymal spermatozoa. (**b**) Track view of H3K27ac ChIP-seq data with biological replicates for each stage of spermatogenesis. ES: embryonic stem cells. (**c**) Heatmap of distal H3K27ac peaks during spermatogenesis by k-means clustering analysis. (**d**) MAnorm analysis of H3K27ac peaks at each transition of spermatogenesis. The genomic distribution of each peak is shown with colored bars.

For the quantitative comparison of active enhancers during spermatogenesis, we analyzed H3K27ac ChIP-seq peaks ±1 kb outside transcription start sites (TSSs); hereafter, we refer to such peaks as ‘distal peaks’ and peaks within ±1 kb as ‘proximal peaks.’ Through the use of MACS2^24^, a program that identifies the significant enrichment of ChIP-seq signals, we detected 11,433 distal H3K27ac ChIP-seq peaks that were present in at least one stage of spermatogenesis (for these analyses, we permitted only distal peaks with a normalized enrichment value of ≥4; see Methods; Supplementary Table 1). The distal peaks were categorized into 9 clusters through k-means clustering; we further organized these into three classes as follows (Fig. 1c). The first class (705 peaks, comprising clusters 1-3) represents constitutive active enhancers, i.e., those observed throughout spermatogenesis. The second class (2,524 peaks, comprising clusters 4 and 5) represents enhancers that are active in the mitotic proliferation phase of spermatogenesis (i.e., the ‘mitotic phase’) but are inactive in meiotic and postmeiotic phases. Notably, the third class (8,204 peaks, comprising clusters 6-10) consists of enhancers that are largely inactive in the mitotic phase yet are highly active in meiotic and postmeiotic stages. The number of H3K27ac ChIP-seq peaks gradually increases over the course of spermatogenic differentiation. Taken together, on the contrary to a previous view that mouse spermatogenesis is not involved in enhancer activation^8^, these results demonstrate that large numbers of active enhancers are established *de novo* in spermatogenesis.

To elucidate how active enhancers change in the course of spermatogenesis, we examined the dynamics of H3K27ac peaks at various transitions from one stage to another. For these analyses, we used MAnorm, a peak-analysis program that facilitates quantitative comparisons of peaks derived from two pairwise next-generation sequencing datasets (see Methods)^25^. Although H3K27ac proximal peaks were largely common at each transition, distal H3K27ac peaks representative of active enhancers underwent dynamic changes in spermatogenesis (Fig. 1d). With regard to distal H3K27ac peaks, a large fraction persisted between THY1^+^ and KIT^+^ spermatogonia; we refer to such peaks as ‘common’ (Fig. 1d). These data suggest that, for the most part, THY1^+^ and KIT^+^ spermatogonia share a largely common profile of active enhancers. In comparison, MAnorm analysis between KIT^+^ spermatogonia and PS revealed a dynamic change in the distribution of active enhancers at the mitosis-to-meiosis transition. This result suggests that a majority of active enhancers in KIT^+^ spermatogonia disappear prior to meiosis, and the extensive *de novo* formation of active enhancers takes place in meiotic prophase (Fig. 1d). For the large part, this *de novo* formation of active enhancers occurred in intergenic and intronic regions. The continued alteration of active enhancers occurred from meiotic PS to postmeiotic RS, and we observed the additional *de novo* establishment of active enhancers in RS (Fig. 1d).

### *De novo* establishment of super-enhancers after the mitosis-to-meiosis transition facilitates robust expression of key spermatogenesis genes

To elucidate how massive, dynamic transcriptional change is stimulated at the mitosis-to-meiosis transition, we sought to test the following hypothesis: The transcriptional change of the mitosis-to-meiosis transition is associated with the establishment of “super-enhancers (SEs).” SEs are genomic regulatory units made up of multiple enhancers, bound by transcription factors to control key regulatory genes required for cellular identity^17,19,26^. SEs have been defined as large chromatin domains enriched with H3K27ac and/or other active enhancer marks^27^; drawing on this definition, we identified SEs based on elevated H3K27ac enrichment in spermatogenesis using the following criteria: 1) Enhancers within 12.5 kb of each other were consolidated into a single entity; (2) enhancer entities were ranked by H3K27ac ChIP-seq signal intensity; finally, (3) enhancer entities enriched with H3K27ac signal above a certain cutoff were defined as SEs (see Methods). Using this definition, we found that SEs are established in the course of spermatogenesis, and they increase in number as germ cells mature: We identified 65 SEs in THY1^+^ spermatogonia, 182 SEs in KIT^+^ spermatogonia, 487 SEs in PS, and 1,114 SEs in RS (Fig. 2a, Supplementary Table 2). The number of SEs established *de novo* increases as spermatogenesis progresses (Fig. 2b). Among the 65 SEs in THY1^+^ spermatogonia, 85% (55/65) are common to SEs identified in KIT^+^ spermatogonia (Fig. 2c), indicating a largely common profile of SEs in mitotically dividing THY1^+^ and KIT^+^ spermatogonia. However, among the 182 SEs in KIT^+^ spermatogonia, only 32% (59/182) are common to SEs in PS (Fig. 2b). These data reveal the dynamic, *de novo* formation of SEs at the mitosis-to-meiosis transition. After the mitosis-to-meiosis transition, 57% (278/487) of SEs in PS were common to SEs in RS; we observed the establishment of 836 new SEs in RS (Fig. 2b).

**Figure 2.**
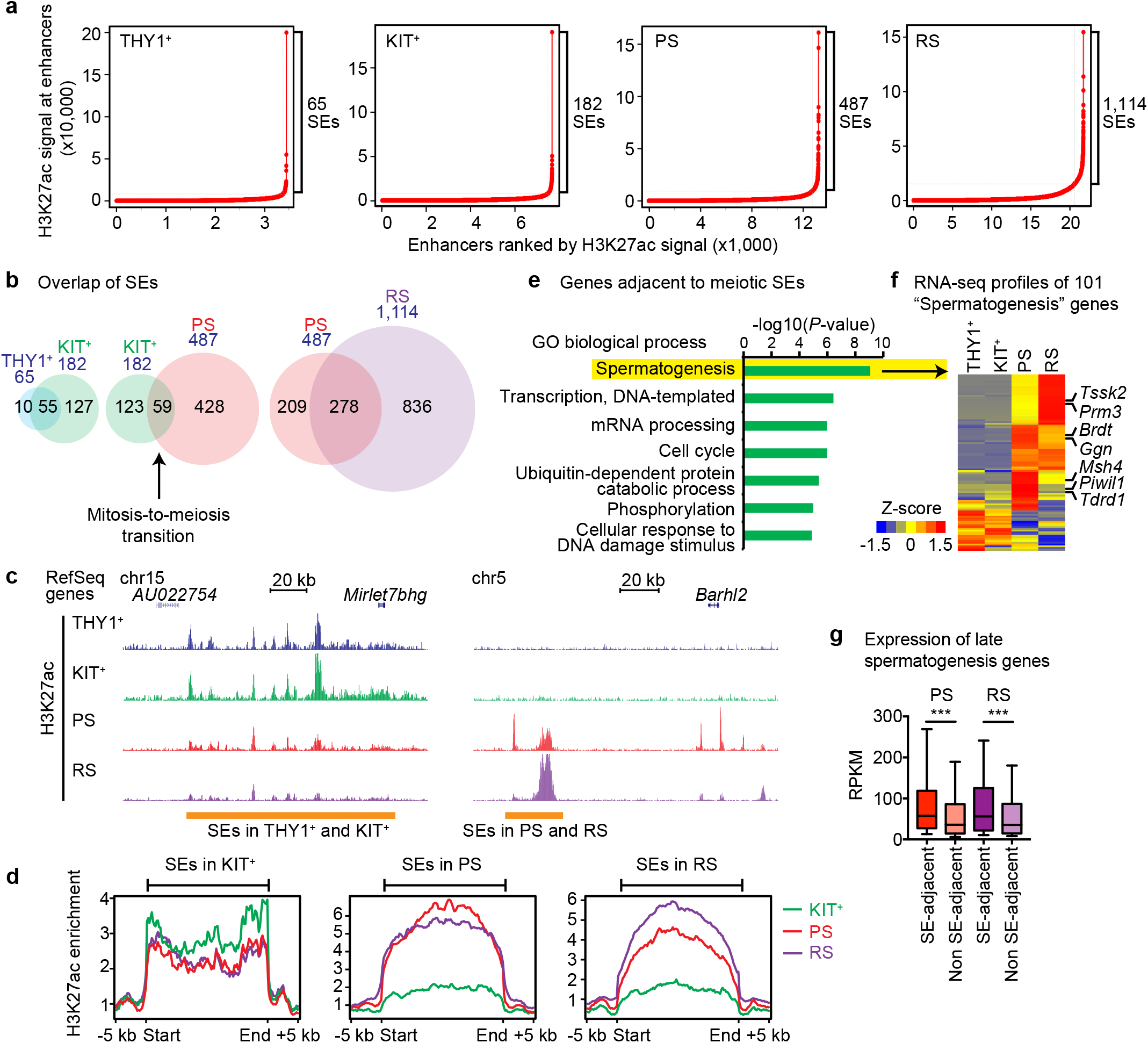
Identification of super-enhancers during spermatogenesis. (**a**) Identification of SEs in representative stages of spermatogenesis. (**b**) Overlap of SEs in each transition during spermatogenesis. (**c**) Track view of H3K27ac ChIP-seq data on representative SEs in spermatogenesis. (**d**) Average tag density of H3K27ac ChIP-seq reads at SEs. (**e**) Gene ontology analysis of genes adjacent to meiotic SEs. (**f**) RNA-seq profiles of 101 “spermatogenesis” genes. (**g**) Box-and-whisker plots showing distribution of RPKM values for RNA-seq data of late spermatogenesis genes. Central bars represent medians, the boxes encompass 50% of the data points, and the whiskers indicate 90% of the data points. *** *P* < 0.0001, Mann-Whitney U test.

We identified distinct characteristics of SEs common to the mitotic stages (i.e., between THY1^+^ and KIT^+^ spermatogonia), and the SEs common to the meiotic (i.e., PS) and postmeiotic (i.e., RS) stages. First, with respect to SEs common to mitotic stages (THY1^+^ and KIT^+^ spermatogonia), H3K27ac was decreased in PS, but tended to persist throughout spermatogenesis into at least as late as the RS stage; for example, H3K27ac was still present after the resolution of a SE on chromosome 15 (Fig. 2c). We designate SEs unique to THY1^+^ and/or KIT^+^ spermatogonia ‘mitotic SEs.’ Through average tag density analyses, we confirmed that decreased H3K27ac persisted to as late as the RS stage at the genomic loci of mitotic SEs (Fig. 2d). On the other hand, with respect to SEs unique to PS, H3K27ac was largely absent from corresponding genomic loci in THY1^+^ and KIT^+^ spermatogonia, but H3K27ac was robustly established during the mitosis-to-meiosis transition (Fig. 2c, d). We term SEs unique to PS and/or RS ‘meiotic SEs.’ Intriguingly, meiotic SEs tend to consist of large and broad H3K27ac peaks, while mitotic SEs tend to comprise clusters of distinct, narrow H3K27ac peaks (Figs. 2c and 2d); SEs with this distinct, narrow conformation have been reported as a general feature of tissue-specific SEs in other mitotically proliferating cells^17–19^.

Next, we examined whether meiotic SEs are associated with specific gene expression programs after the mitosis-to-meiosis transition. Gene ontology analysis revealed that genes adjacent to meiotic SEs (i.e., genes within 20 kb upstream to 50 kb downstream of SEs) are enriched for roles in “spermatogenesis” and “male gamete function” (Fig. 2e). We identified 101 genes that were categorized for “spermatogenesis,” and this gene group is highly expressed after the mitosis-to-meiosis transition (Fig. 2f, Supplementary Table 3). This group includes key regulators of spermatogenesis such as *Brdt,* a key chromatin regulator in spermatogenesis^28^; and *Trdr1* and *Piwil1* (also known as *Miwi*), components of the Piwi-interacting RNA pathway^29,30^; *Msh4,* an essential gene for meiotic recombination^31^; and other spermatogenesis genes including *Ggn, Prm3,* and *Tssk2* (Fig. 2f). Next, we investigated whether genes adjacent to meiotic SEs are subject to higher expression than genes that are not adjacent to meiotic SEs. Among 2,623 late spermatogenesis genes (i.e., genes that are not highly expressed in spermatogonia but are highly expressed in PS and/or RS by a ≥4-fold change compared to spermatogonia: a gene list is included in Supplementary Table 4), 652 genes are located adjacent to meiotic SEs (i.e., genes within 20 kb upstream to 50 kb downstream of SEs). These genes are robustly expressed compared to the remaining 1,971 late spermatogenesis genes that are not adjacent to SEs (Fig. 2g). These results suggest that the *de novo* establishment of meiotic SEs facilitates robust expression of key spermatogenesis genes.

### A-MYB binding is associated with the establishment of meiotic SEs for targeted activation of germline genes

Enhancers contain transcription factor (TF)-binding sites to regulate the expression of target genes^32^. Thus, we sought to identify TF-binding motifs that underlie gene expression programs unique to spermatogenesis, specifically those associated with the mitosis-to-meiosis transition. Using the motif analysis program MEME-ChIP^33^, we identified consensus motifs similar to known TF-binding motifs in active enhancers. To identify TF-binding sites indicative of the mitotic stage versus the meiotic stage, we compared H3K27ac ChIP-seq peaks between KIT^+^ spermatogonia versus PS. Among the H3K27ac peaks unique to KIT^+^, the TF-binding motif with the lowest *E* value (i.e., an *E* value = 4.3×10^-16^; *E* values are expected values output by the MEME expectation maximization algorithm)^34^ contained motifs for STAT family transcription factors (STAT1, STAT3, and STAT5a). This is in line with the function of STAT3 in spermatogonial differentiation^35^. The motif with the second lowest *E*-value (*E*-value = 1.5×10^-13^) contains a common binding motif for FOX family transcription factors (FOXK1, FOXJ3, and FOXG1).

Of note, these data for motif enrichment in KIT^+^ spermatogonia are distinct from data obtained for KIT^+^ spermatogonia in our previous study of accessible chromatin (detected via ATAC-seq; like H3K27ac ChIP-seq speaks, ATAC-seq peaks are indicators of cis-regulatory elements)^9^. To more carefully examine the union and/or exclusivity of motifs in H3K27ac and ATAC peaks, we performed HOMER motif analyses^36^ using an expanded TF binding motif library taken from the Cis-BP database^37^ (see Methods). H3K27ac and ATAC double-positive peaks in KIT^+^ spermatogonia contain consensus motifs such as NR5A2, a TF implicated in germ cell development; the retinoid receptors RXRA and RXRB; and binding motifs for FOS, FOSL2, and JUND. These data are common with previously identified motifs in ATAC peaks in KIT^+^ spermatogonia^9^ (Fig. 3b, Supplementary Table 5). Notably, in KIT^+^ spermatogonia, consensus motifs for DMRT1, a key TF that regulates the mitosis-to-meiosis transition^38^, were enriched only in ATAC-positive/H3K27ac-negative peaks, suggesting that DMRT1 functions outside of active enhancers (Fig. 3b). A similar feature was also found in PS: The binding motifs for the POU/OCT family of TFs (POU2F, POU3F, and POU1F1) were found only in ATAC-positive/H3K27ac-negative peaks, suggesting that the POU/OCT family of TFs functions outside of active enhancers in PS (Fig. 3b).

**Figure 3.**
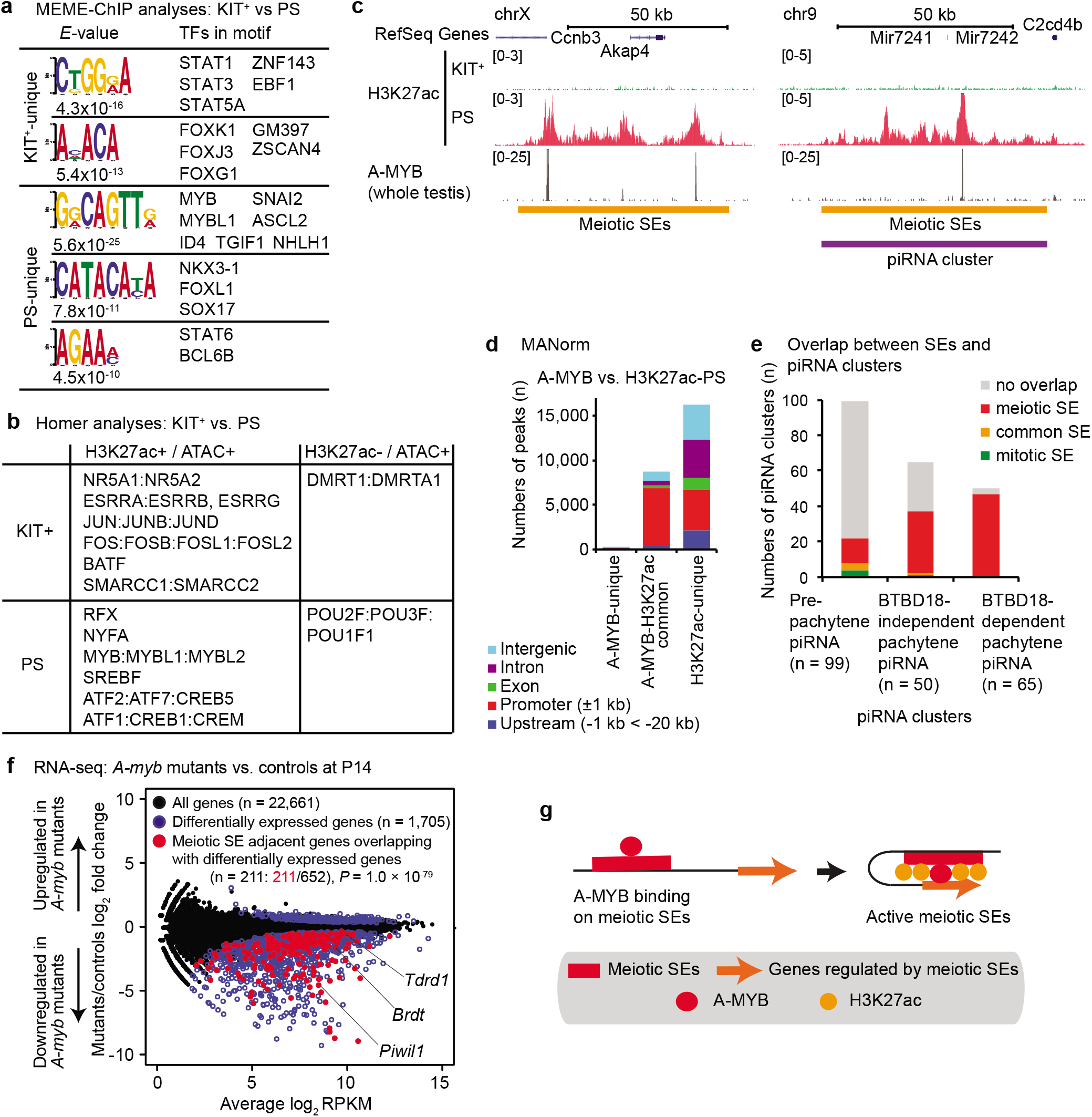
A-MYB binding is associated with the establishment of meiotic SEs for targeted activation of germline genes. (**a**) Representative data of MEME-ChIP analysis of H3K27ac reads between KIT^+^ and PS. (**b**) Summary of HOMER analysis of H3K27ac reads between KIT^+^ and PS. (**c**) Track view of H3K27ac ChIP-seq data on meiotic SEs in spermatogenesis. (**d**) MAnorm analysis between A-MYB peaks in testes and PS-H3K27ac peaks. (**e**) Overlap between SEs and piRNA clusters. (**f**) RNA-seq analysis of *A-myb* mutant versus heterozygous control testes at 14 days post-partum (dpp). The 1,705 genes evincing significant changes in expression (>2-fold change, *P adj* < 0.01: a binomial test) in *A-myb* mutants are represented by blue circles. *P* value is based on a hypergeometric probability test. The 211 dysregulated genes (represented by red circles)/652 meiotic SE-adjacent genes (*P* = 1.0 × 10^-79^) compared to 1,705 dysregulated genes/all 22,661 RefSeq genes in the genome. (**g**) A model of A-MYB dependent establishment of meiotic SEs.

The PS-unique motifs found within H3K27ac peaks revealed a common and notable feature between MEME-ChIP and HOMER motif analyses: A strong enrichment of MYB binding motifs was identified by both analysis pipelines (MEME-ChIP analysis, GGCAGTT: *E* value = 5.6×10^-25^, Fig. 3a; HOMER motif analysis, Fig. 3b, Supplementary Table 5). These MYB motifs are recognized by A-MYB (also known as MYBL1), a key transcription factor for germline genes^20^, and are also strongly enriched in A-MYB testes ChIP-seq peaks^21^. Using this previously published dataset, we identified A-MYB ChIP-seq peaks at the center of SE-associated H3K27ac peaks on the X chromosome and autosomes in PS (Fig. 3c). Given the role of A-MYB in the regulation of meiotic transcription, these data raise the possibility that A-MYB binding may nucleate establishment of meiotic enhancers and SEs onto the surrounding chromatin. MAnorm analyses revealed that, genome-wide, most A-MYB peaks overlap H3K27ac peaks (Fig. 3d), which further supports the function of A-MYB in priming meiotic enhancers.

Importantly, we found that meiotic SEs overlap pachytene piRNA clusters, which produce pachytene piRNAs (Fig. 3c, right panel, and Fig. 3e). Among pachytene piRNA clusters, we found that BTBD18-dependent piRNA loci are highly likely to overlap meiotic SEs (Fig. 3e); BTBD18 is an essential factor for pachytene piRNA production by way of transcriptional elongation^39^. Since A-MYB is essential for the production of pachytene piRNA^21^, these data lend further support to the assertion that A-MYB functions in the priming of meiotic SEs; furthermore, A-MYB and meiotic SEs may comprise, in part or whole, a potential mechanism for the production of pachytene piRNA.

To test whether meiotic SE-associated genes are regulated by A-MYB, we analyzed previously published RNA-seq data from the testes of *A-myb* mutants (*Mybl1^repro9^*) and heterozygous littermate controls at P14^35^ (Fig. 3f). We observed a significant overlap of meiotic SE adjacent genes and genes differentially expressed in *A-myb* mutants (211 differentially expressed genes out of 652 meiotic SE adjacent genes (32.7%); this is in comparison to 1,705 differentially expressed genes out of all 22,661 RefSeq genes in the genome (7.5%); *P* = 1.0×10^-79^; hypergeometric probability test), and many of the differentially expressed genes were found in the downregulated genes of *A-myb* mutants. Together, these results suggest that A-MYB-binding might trigger the establishment of meiotic SEs to activate target germline genes (Fig. 3g).

### SCML2 is required for the resolution of mitotic SEs during meiosis

Next, we sought to determine a mechanism underlying the resolution of mitotic SEs at the mitosis-to-meiosis transition. We focused our investigation on the function of SCML2, a germline specific Polycomb protein that is responsible for dynamic transcriptional changes at the transition^3^; in mice deficient for *Scml2* (i.e., *Scml2*-knockout (KO) mice), somatic/progenitor genes were derepressed on autosomes after the mitosis-to-meiosis transition, and robust activation of late-spermatogenesis genes was compromised as well^3^. Although H3K27ac peaks were comparable between the THY1^+^ and KIT^+^ spermatogonia of both *Scml2*-KO mice and wild-type littermate controls (Supplementary Fig. 2a), MAnorm analysis revealed a large number of unique H3K27ac peaks at intergenic and intronic regions in PS of *Scml2*-KO mice compared to wild-type controls (at intergenic regions, 1,549 peaks in wild-type and 1,951 peaks in *Scml2*-KO; at intronic regions, 845 peaks in wild-type and 4,461 peaks *Scml2*-KO; Fig. 4a). Intriguingly, the increased number of H3K27ac peaks in *Scml2*-KO PS appeared to result from the retention of mitotic enhancers after the mitosis-to-meiosis transition: H3K27ac peaks at mitotic SEs, which are resolved in wild-type PS, were, for the most part, retained in *Scml2*-KO PS and RS (Supplementary Fig. 2b). Average tag density analysis confirmed the genome-wide retention of H3K27ac at SEs from *Scml2*-KO KIT+ spermatogonia to PS and RS (Fig. 4b). Given this evidence, and since SCML2 suppresses somatic/progenitor genes in meiosis^3^, these results suggest that the SCML2-mediated resolution of mitotic SEs constitutes a potential mechanism for the suppression of somatic/progenitor genes at the mitosis-to-meiosis transition.

**Figure 4.**
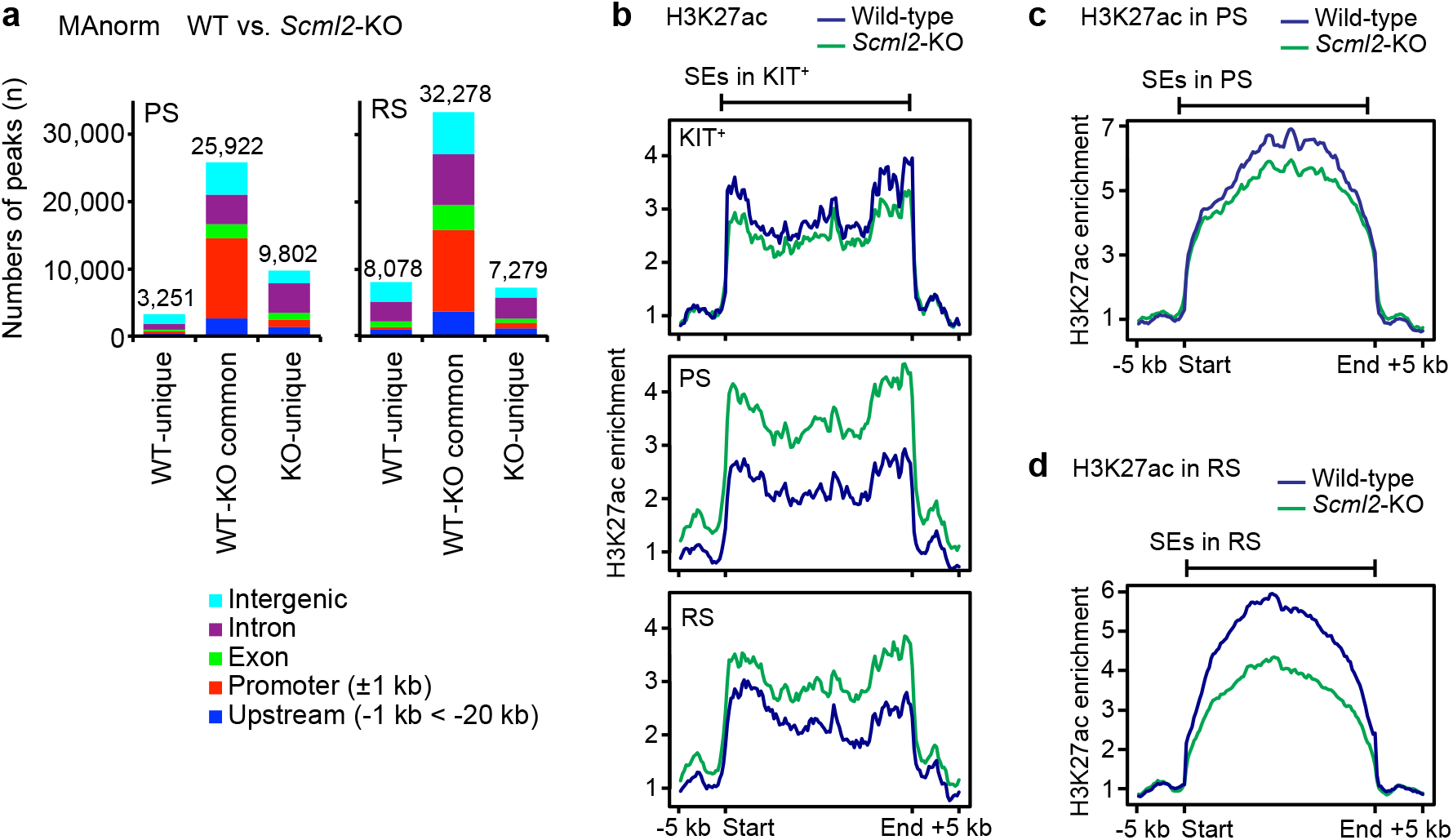
SCML2 is required for the resolution of mitotic SEs during meiosis. (**a**) MAnorm analysis of H3K27ac peaks in PS and RS between wild-type and *Scml2*-KO. (**b**) Average tag density of H3K27ac ChIP-seq reads at SEs in KIT^+^ in each stage of spermatogenesis (KIT^+^, PS, RS). (**c, d**) Average tag density of H3K27ac ChIP-seq reads at SEs in PS (**c**) and SE in RS (**d**).

We also observed that the intensity of H3K27ac at meiotic SEs was slightly decreased in *Scml2*-KO PS (Fig. 4c), and *Scml2*-KO RS saw a further decrease in H3K27ac intensity (Fig. 4d). These observations are consistent with the down-regulation of late-spermatogenesis genes in PS and RS *Scml2*-KO mice^3^. SCML2 is required for the establishment of H3K27me3 during meiosis, forming two major classes of bivalent genomic domains comprised of H3K27me3 and H3K4me2/3: Class I domains, which are associated with developmental regulator genes; and Class II domains, which are associated with somatic/progenitor genes^40^. We observed an increase in H3K27ac signal intensity at both classes of bivalent domains in *Scml2*-KO mice (Supplementary Fig. 2c). We presume that this is the consequence—at least in part—of an antagonistic relationship between H3K27me3 and H3K27ac, since both post-translational modifications occupy the same amino acid residue (K27) of the histone H3 tail.

### SCML2 is required for the formation of SEs on the X chromosome during meiosis

Switching focus to meiosis, we performed analyses to elucidate the mechanisms governing the establishment of active enhancers on the male sex chromosomes. During male meiosis, the sex chromosomes undergo regulation distinct from autosomes due to a central regulatory mechanism known as meiotic sex chromosomes inactivation (MSCI)^41,42^. MSCI engages a DNA damage response (DDR) pathway to catalyze and regulate sex chromosome gene silencing in PS, an essential occurrence prior to the activation of a subset of sex chromosome genes in RS^43,44^. RNF8, an E3 ligase and key DDR factor, is responsible for the establishment of ubiquitination on the sex chromosomes, along with the establishment of active histone modifications such as the enhancer mark H3K27ac, thereby regulating the activation of a subset of sex chromosome genes that escape post-meiotic silencing^23,44^. Contrasting with our genome-wide profiles of H3K27ac (Fig. 1), we observed a scarcity of distal H3K27ac peaks on the sex chromosomes of THY1^+^ and KIT^+^ spermatogonia (Fig. 5a). Of note, in accordance with the chromosome-wide establishment of H3K27ac signals on the XY body as detected by fluorescence microscopy^23^, many H3K27ac peaks (930 intergenic and 327 intronic peaks of total 1851 peaks) were established *de novo* on the sex chromosomes in the intergenic and intronic regions from KIT+ spermatogonia to PS (Fig. 5a).

**Figure 5.**
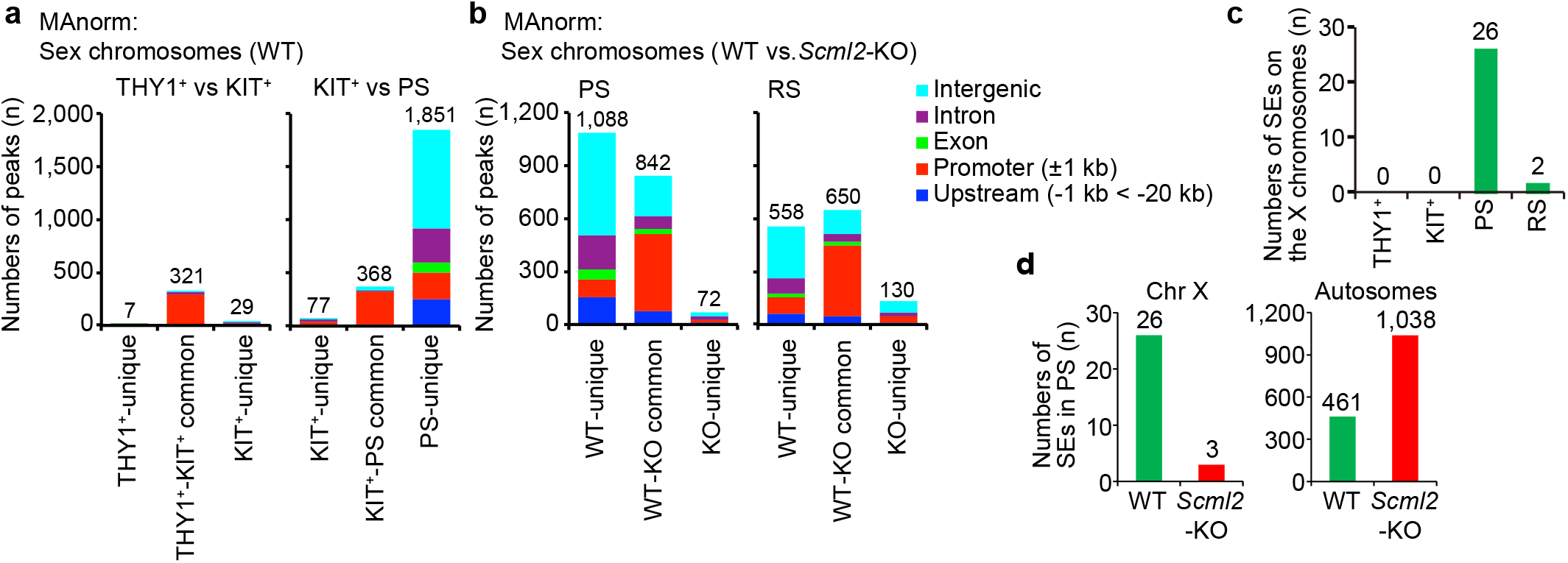
SCML2 is required for the formation of SEs on the X chromosome during meiosis. (**a**) MAnorm analysis of H3K27ac peaks on the sex chromosomes between THY1^+^ and KIT^+^ spermatogonia, and between KIT^+^ spermatogonia and PS. (**b**) MAnorm analysis of H3K27ac peaks on the sex chromosomes in PS and RS between wild-type and *Scml2*-KO. (**c**) Numbers of SEs on the X chromosome in each stage of spermatogenesis. (**d**) Number of SEs in PS on the X chromosome and autosomes.

To dissect the regulatory mechanisms underlying this process, we focused on SCML2, which has a critical regulatory function on the sex chromosomes independent of its functions on autosomes^3^. SCML2 functions downstream of the DDR pathway that initiates MSCI, where it cooperates with RNF8 to establish H3K27ac^23^. MAnorm analysis revealed that, on the sex chromosomes of PS and RS, a large portion of distal H3K27ac peaks (particularly those in intergenic and intronic regions) depend on SCML2 (Fig. 5b). Our prior report showed that, interestingly, ATAC-seq peaks appeared specifically on the sex chromosomes of PS in an SCML2-dependent fashion too^9^. Together, these data indicate that SCML2 is a key regulatory factor for chromatin accessibility and H3K27ac deposition on the sex chromosomes in meiosis. Accordingly, 26 SEs are established on the X chromosomes in meiosis (Fig. 5c), and these largely depend on SCML2 (Fig. 5d). Intriguingly, this is unlike SCML2’s function to resolve mitotic SEs after the mitosis-to-meiosis transition (Fig. 4); as such, we observed increased numbers of SEs on the autosomes of *Scml2*-KO PS (Fig. 5d). Together, these results demonstrate distinct autosome- and sex chromosome-specific functions for SCML2 in the regulation of enhancers in spermatogenesis.

### Meiotic super-enhancers on autosomes are poised in undifferentiated spermatogonia

SEs on the sex chromosomes are established downstream of the DDR pathway that initiates MSCI. This DDR-dependent regulation of the sex chromosomes is specific to the unpaired — or unsynapsed — status of the hemizygous male sex chromosomes in meiosis, when homologous autosomes otherwise fully synapse to facilitate recombination^42^. Given this difference between sex chromosomes and autosomes, we suspected that SEs on the autosomes are regulated by a distinct mechanism. Thus, to determine the mechanism by which autosomal SEs are established, we examined the epigenetic status of meiotic SEs in progenitor cells. We examined H3K4me2 and H3K4me3, active marks that were previously reported to be associated with poised gene promoters during spermatogenesis^4^. Notably, prior to the establishment of H3K27ac, H3K4me2 was present on autosomal meiotic SEs in THY1^+^ and KIT^+^ spermatogonia (Fig. 6a). Additionally, H3K4me3 is also enriched on autosomal meiotic SEs in THY1^+^ spermatogonia (Fig. 6a). These features were unique to meiotic SEs: Other meiotic enhancers detected through analyses of distal H3K27ac peaks did not exhibit these features (Supplementary Fig. 3a). These results suggest that meiotic SEs are poised as early as the THY1^+^ spermatogonia phase to prepare for the expression of key spermatogenesis genes after the mitosis-to-meiosis transition. These features were not observed on meiotic SEs associated with the X chromosome (Fig. 6b), lending further support for the distinct regulation of meiotic SEs between autosomes and the X chromosome. While poised chromatin is unique to autosomal meiotic SEs—and not associated with other meiotic enhancers—we found that the TSSs of late spermatogenesis genes are also broadly poised for activation in spermatogonia (Supplementary Fig. 3b)^4^. Together, these data indicate that SE-associated late spermatogenesis genes on autosomes are poised for activation in two layers: SEs and TSSs (Fig. 6c). We propose that this form of epigenomic programming ensures the unidirectional differentiation of spermatogenesis (Fig. 6c).

**Figure 6.**
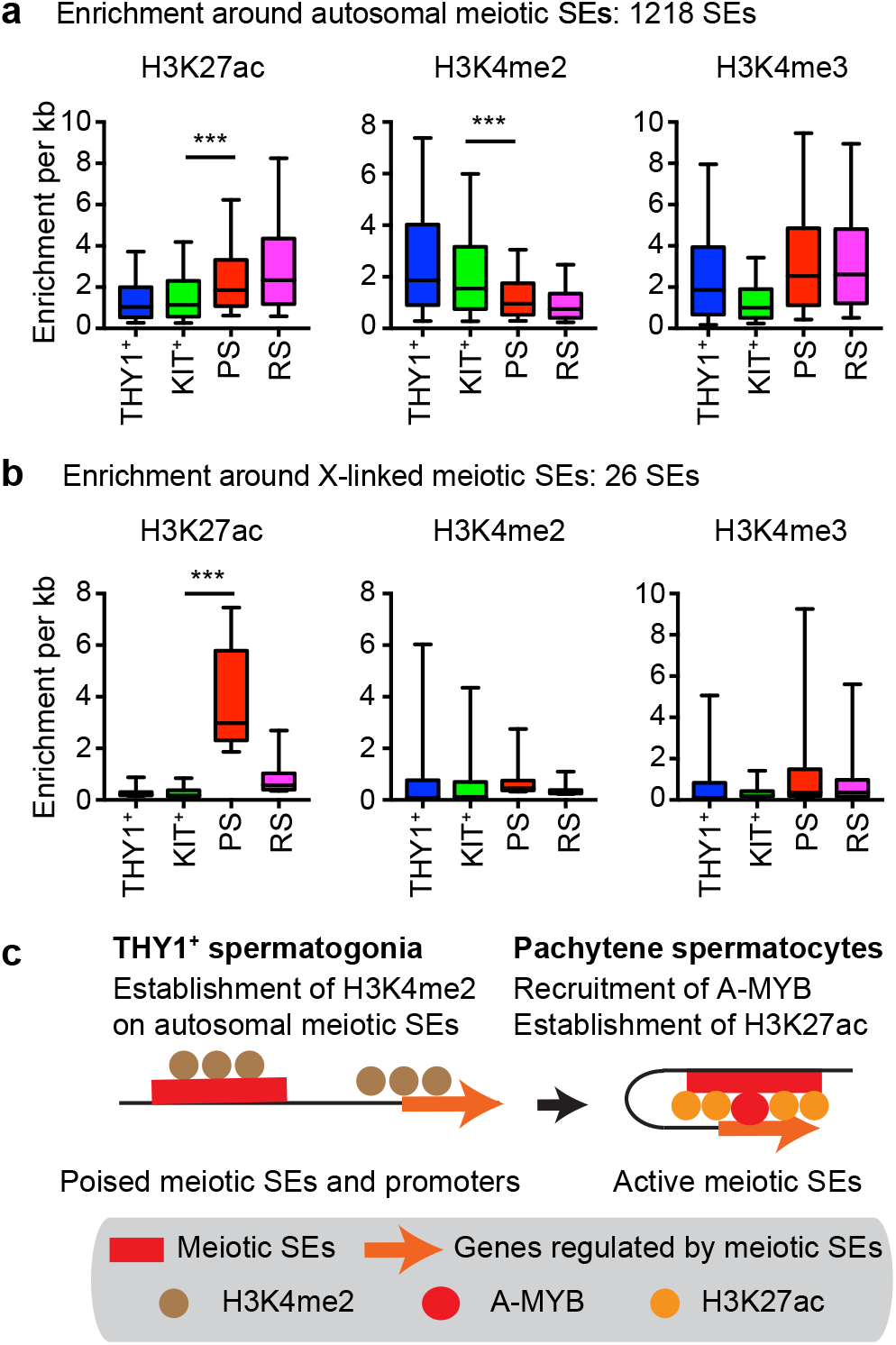
Meiotic super-enhancers on autosomes are poised in undifferentiated spermatogonia. (**a, b**) Box-and-whisker plots showing distribution of ChIP-seq read enrichment around autosomal meiotic SEs (**a**) and around X-linked meiotic SEs (**b**). Central bars represent medians, the boxes encompass 50% of the data points, and the whiskers indicate 90% of the data points. *** *P* < 0.0001, Mann-Whitney U test. (**c**) A model of poised meiotic SEs and promoters in THY1+ spermatogonia and establishment of active meiotic SEs in pachytene spermatocytes.

Next, we sought to identify mechanisms for the expression of postmeitoic spermatid specific genes. On autosomes in PS, at distal H3K27ac peaks around RS-specific genes, we observed an increase in H3K27ac and H3K4me3 signals prior to activation in RS, and H3K27ac and H3K4me3 signals became highly enriched in RS (Supplementary Fig. 3c). On the other hand, on the X chromosome in PS, at distal H3K27ac peaks around RS-specific genes, H3K27ac and H3K4me2 became temporary enriched in PS prior to gene activation in RS (Supplementary Fig. 3d). Since, on the sex chromosomes, accumulation of H3K4me2 and H3K27ac takes place downstream of RNF8 function, and accumulation of H3K4me2 and H3K27ac is regulated by SCML2 too^23,44^, these results serve to further reveal gene activation mechanisms that are distinct between autosomes and the sex chromosomes in haploid RS.

### Identification of key regulatory factors for both mitotic and meiotic enhancers

Finally, we took advantage of our new data sets to infer general mechanisms underlying the regulation of mitotic and meiotic enhancers. Since the meiotic gene program is, by and large, repressed in cell types that undergo mitotic divisions, we sought to identify putative TFs that meet one of two counteractive conditions: (1) Those that can operate on and/or promote the activity of mitotic enhancers, and (2) those that can suppress meiotic enhancers. To this end, we used our recently published RELI (Regulatory Element Locus Intersection) algorithm^45^, to compare the genomic locations of our H3K27 peak data with a large collection of publicly available ChIP-seq data. Taking the genomic location information for H3K27ac peaks detected to be mitotic enhancers in KIT^+^ spermatogonia, we analyzed the intersections between these data and publicly available ChIP-seq data sets for many TFs in many contexts. Since the overwhelming majority of public ChIP-seq data are from somatic cells that undergo mitotic divisions in between cell cycles, this informs (at least) one interpretation for such an experiment: The enrichment of intersections between mitotic enhancers and TFs could be indicative of general mechanisms that operate on mitotic enhancers. In addition to TFs that were previously associated with spermatogonia, such as STAT3, TCF3, MAZ, and ETS1^35,46,47^, we identified additional factors with enriched ChIP-seq peaks at distal H3K27ac peaks in KIT^+^ spermatogonia: SRF, TCF12, GATA4, BCL6, CEBPB, and MAX (Fig. 7a, Supplementary Table 6). When we applied the same analysis to mitotic SEs, we identified UBTF, RBP1, CHD1, ZFX, and KLF4 as specific factors that may be involved in their regulation (Fig. 7b, Supplementary Table 7). Among them, ZFX was previously implicated in spermatogenesis^48^. Next, we applied this strategy to identify factors that suppress meiotic enhancers in the mitotic phase. Compellingly, at the sites for meiotic enhancers as determined by the loci of distal H3K27ac peaks in PS, we revealed high enrichment for factors that comprise, in part, transcriptional silencing machinery, including REST, TRIM28, RCOR2, SIN3A, and YY1 (Fig. 7c, Supplementary Table 8). Of note, when we applied this analysis to meiotic SEs in the mitotic phase, we identified KDM5A, a histone demethylase that acts on H3K4me3, as the factor having the highest enrichment at meiotic SEs (Fig. 7d, Supplementary Table 9). Together, these analyses systematically identify putative regulators of mitotic and meiotic enhancers, providing clues for understanding their underlying molecular mechanisms.

**Figure 7.**
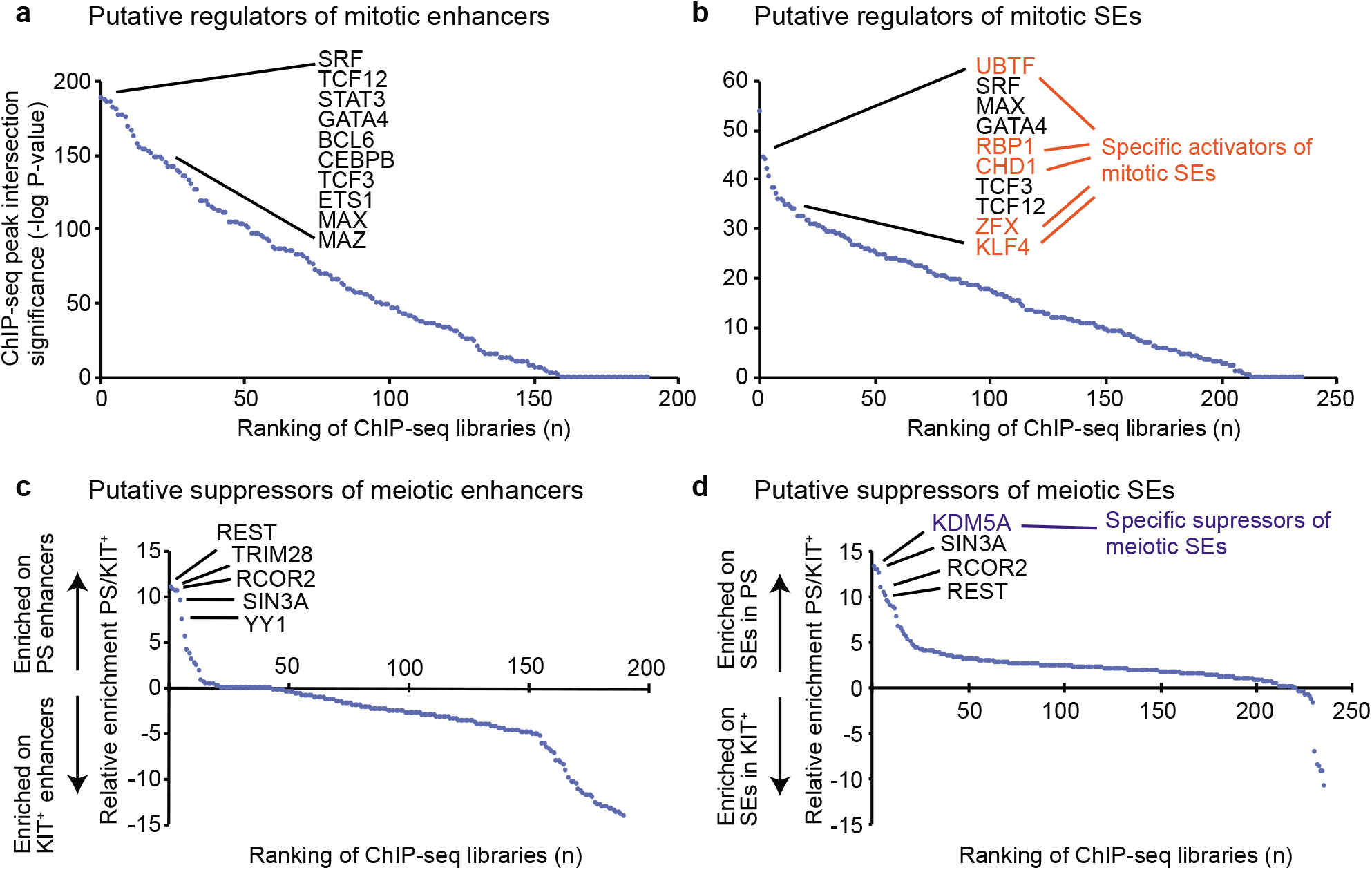
Identification of key regulatory factors for mitotic and meiotic enhancers and SEs. (**a, b**) Identification of putative regulators for mitotic enhancers and SEs. The Y-axis indicates the –log of the P-value for the overlap between publically available ChIP-seq datasets for various TFs and mitotic enhancers (**a**) or SEs (**b**) based on the RELI algorithm (see Methods). TFs of interest are highlighted. (c, d) Comparison between enriched TFs in meiotic PS vs. KIT^+^ enhancers (**c**) or SEs (**d**). The Y-axis indicates the ratio of the–log of the P-value for the overlap between publically available ChIP-seq datasets for various TFs and enhancers (**c**) or SEs (**d**) based on the RELI algorithm.

## Discussion

In this study, we determined the profiles of active enhancers in representative stages of spermatogenesis, and we demonstrated that SE switching underlies the dynamic transcriptome change of the mitosis-to-meiosis transition. Our results establish an overarching molecular logic for this switching: SCML2 resolves mitotic SEs, and A-MYB establishes meiotic SEs to regulate meiotic transcription. Recent reports indicate that it is, generally speaking, the nature of SEs to regulate gene expression that underlies the cellular identity of cell types^17,18^; with this in mind, the A-MYB-dependent regulation of meiotic SEs becomes a conceivable mechanism for gene expression that defines the cellular identity of germ cells in late spermatogenesis. Our analyses revealed SE-adjacent genes such as *Brdt, Miwil1,* and *Tdrd1*—genes that are critical for late spermatogenesis. Thus, these genes may be candidate genes for cellular identity. In particular, *Miwil1* and *Tdrd1* are involved in the regulation of piRNA^29,30^, and A-MYB is required for the expression of *Miwil1* and *Tdrd1*^21^. For example, A-MYB directly regulates transcription of pachytene piRNA clusters, and robust pachytene piRNA production is thought to be regulated by a feed forward loop as part of the transcriptional activation of piRNA clusters and the expression of piRNA regulators such as MIWIL1 and TDRD1^21^. Our data support the possibility that this feedforward loop is mediated by the establishment of meiotic SEs. Interestingly, the A-MYB-dependent activation of germline genes is an ancient mechanism also found in rooster testes^21^. So, it is interesting to speculate that the establishment of meiotic SEs by A-MYB could be an ancient mechanism too—and one that was possibly exploited by, or otherwise coopted by, pachytene piRNA production in the course of evolutionary history. Given the robust and evolutionarily conserved nature of germline gene activation via SEs, such a mechanism stands in stark contrast to a concomitant mechanism whereby rapidly evolved enhancers, driven by endogenous retroviruses, activate species-specific genes after the mitosis-to-meiosis transition of spermatogenesis (Sakashita et al., co-submitted).

In tumor cells, SEs are regulated by BRD4, a member of the bromodomain and extraterminal (BET) subfamily of proteins, and inhibition of BRD4 results in the dysregulation of SE-associated genes, including the *MYC* proto-oncogene^26^. In spermatogenesis, a testis-specific member of the BET family, BRDT, is required for the meiotic gene expression program^28^, and small-molecule inhibition of BRDT caused spermatogenic failure^49^. Given the molecular similarities between BRD4 and BRDT, it is possible that BRDT could be a binding protein for meiotic SEs, and loss of function of BRDT could represent loss of function of meiotic SEs. Since we identified *Brdt* as a meiotic SE-adjacent gene, BRDT may contribute to a feedforward loop that putatively establishes meiotic SEs. Curiously, another protein containing bromodomains, BRWD1 (Bromodomain And WD Repeat Domain Containing 1), can also recognize acetylated lysine residues and is required for postmeiotic transcription in spermatids^50^. Likewise, BRD4 is also associated with gene expression in spermatids^51^. So, given the increasing numbers of active enhancers established as PS progresses to become RS (Fig. 1), these observations collectively represent an important direction for future work: to determine the functions of bromodomain proteins in the regulation of germline enhancer activity.

Our study elucidated distinct forms of regulation for active enhancers on autosomes versus active enhancers on sex chromosomes in spermatogenesis; these forms of regulation are mediated by the distinct functions of SCML2 on autosomes versus sex chromosomes. On the autosomes, SCML2, a highly expressed protein in undifferentiated spermatogonia^3^, is involved in the resolution of mitotic SEs after the mitosis-to-meiosis transition (Fig. 4), while meiotic SEs are already poised with H3K4me2 in undifferentiated spermatogonia (Fig. 6). Therefore, it is conceivable that these dual mechanisms preprogram meiotic gene expression as early as the undifferentiated spermatogonia phase of spermatogenesis, all in preparation for the unidirectional differentiation of spermatogenesis. On the other hand, on the sex chromosomes, H3K27ac deposition depends on RNF8^23^, a DDR factor, in addition to SCML2 (Fig. 5). Given that MDC1, a DDR factor and key regulator of MSCI^43^, is necessary for the localization of SCML2 on the XY body^3^, our results indicate that active enhancers and postmeiotic gene expression are directly downstream of a DDR pathway specific to the sex chromosomes.

Finally, through genome-wide analyses using publicly available ChIP-seq data from many different cell types, we revealed transcription factors that might bind mitotic and meiotic enhancers, as well as mitotic and meiotic SEs (Fig. 7). Among the factors we identified, the transcriptional repressor KDM5A (also known as RBP2 or JARID1A) evinced the highest enrichment value for meiotic SEs. This is of particular interest because KDM5A was originally identified as an RB (Retinoblastoma)-binding protein and is implicated in tumorigenesis^52^. Since many germline-associated genes are expressed in many cancer types—or, put another way, many germline-associated genes are so-called cancer/testis genes^53^—it is interesting to consider that the regulation of meiotic SEs could, in turn, drive or otherwise regulate germline gene expression in various cancers.

In summary, our current study provides a framework to understand enhancer activity and the regulation of gene expression during spermatogenesis. Because our study focuses on representative stages, it will be important to further dissect these mechanisms in order to fully understand the complex and well-coordinated nature of spermatogenesis. Recent studies using single cell analyses have revealed new details on the shifting, transitory transcriptomic and epigenomic environments of progressive cell types in human and mouse spermatogenesis^54–59^. Such dynamism is achievable through the functional interplay of complex combinations of TFs and enhancers, as well as other regulatory elements. Indeed, more than a thousand TFs are differentially expressed during the mitosis-to-meiosis transition in spermatogenesis^9^. Of note, the testis has the largest number of specifically expressed TFs of all organs^60^. The systematic determination of germline cis-regulatory elements makes for a compelling future research direction to understand germline mechanisms, including fundamental aspects of the mitotic and meiotic programs.

## Methods

### Animals

*Scml2*-KO mice were previously reported ^3^.

### Germ cell fractionation

Pachytene spermatocytes and round spermatids were isolated via BSA gravity sedimentation as previously described^61^. Purity was confirmed by nuclear staining with Hoechst 33342 using fluorescence microscopy. In keeping with previous studies from the Namekawa lab^3,9,11,40^, ≥90% purity was confirmed for all purifications. Spermatogonia were isolated as described previously^8^ and collected from C57BL/6N mice aged 6-8 days. Testes were collected in a 24-well plate in Dulbecco’s Modified Eagle Medium (DMEM) supplemented with GlutaMax (Thermo Fisher Scientific), non-essential amino acids (NEAA) (Thermo Fisher Scientific), and penicillin and streptomycin (Thermo Fisher Scientific). After removing the *tunica albuginea* membrane, testes were digested with collagenase (1 mg/ml) at 34°C for 20 min to remove interstitial cells, then centrifuged at 188×g for 5 min. Tubules were washed with the medium and then digested with trypsin (2.5 mg/ml) at 34°C for 20 min to obtain a single cell suspension. Cells were filtered with a 40-μm strainer to remove Sertoli cells, and the cell suspension was plated in a 24-well plate for 1 h in the medium supplemented with 10% fetal bovine serum, which promotes adhesion of remaining somatic cells. Cells were washed with magnetic cell-sorting (MACS) buffer (PBS supplemented with 0.5% BSA and 5 mM EDTA) and incubated with CD117 (KIT) MicroBeads (Miltenyi Biotec) on ice for 20 min. Cells were washed and resuspended with MACS buffer, and filtered with a 40-μm strainer. Cells were separated by autoMACS Pro Separator (Miltenyi Biotec) with the program “possel.” Cells in the flow-through fraction were washed with MACS buffer and incubated with CD90.2 (THY1) MicroBeads (Miltenyi Biotec) on ice for 20 min. Cells were washed and resuspended with MACS buffer and filtered with a 40-μm strainer. Cells were separated by autoMACS Pro Separator (Miltenyi Biotec) with the program “posseld.” Purity was confirmed by immunostaining.

### ChIP-seq library preparation and sequencing

Cells were suspended in chilled 1× PBS. One-eleventh volume of crosslinking solution (50 mM HEPES-NaOH pH 7.9, 100 mM NaCl, 1 mM EDTA, 0.5 mM EGTA, and 8.8% formaldehyde) was added to the cell suspension and incubated on ice for 8 min. One-twentieth volume of 2 M glycine was added to the cell suspension and incubated at room temperature for 5 min to stop the reaction. Cells were washed twice with PBS, frozen at −80°C, and lysed at 4°C for 10 min each in ChIP lysis buffer 1 (50 mM HEPES pH 7.9, 140 mM NaCl, 10% glycerol, 0.5% IGEPAL-630, 0.25% Triton X-100). After centrifugation at 2,000×*g* for 10 min at 4°C, pellets were resuspended with ChIP lysis buffer 2 (10 mM Tris-HCl pH 8.0, 200 mM NaCl, 1 mM EDTA, 0.5 mM EGTA) and incubated at 4°C for 10 min. After centrifugation at 2,000×g for 10 min at 4°C, pellets were washed with TE containing 0.1% SDS and protease inhibitors (Sigma; 11836145001), and resuspended with the same buffer. Chromatin was sheared to approximately 200–500 bp by sonication using a Covaris sonicator at 10% duty cycle, 105 pulse intensity, 200 burst for 2 min. Sheared chromatin was cleared by centrifugation at 20,000×*g* for 20 min, followed by pre-incubation with Dynabeads Protein G (Thermo Fisher Scientific). Chromatin immunoprecipitation was carried out on an SX-8X IP-STAR compact automated system (Diagenode). Briefly, Dynabeads Protein G were preincubated with 0.1% BSA for 2 h. Then, the cleared chromatin was incubated with beads conjugated to antibodies against H3K27ac (Active Motif; 39133) at 4°C for 8 h, washed sequentially with wash buffer 1 (50 mM Tris-HCl pH 8.0, 150 mM NaCl, 1 mM EDTA, 0.1% SDS, 0.1% NaDOC, and 1% Triton X-100), wash buffer 2 (50 mM Tris-HCl pH 8.0, 250 mM NaCl, 1 mM EDTA, 0.1% SDS, 0.1% NaDOC, and 1% Triton X-100), wash buffer 3 (10 mM Tris-HCl pH 8.0, 250 mM LiCl, 1 mM EDTA, 0.5% NaDOC, and 0.5% NP-40), wash buffer 4 (10 mM Tris-HCl pH 8.0, 1 mM EDTA, and 0.2% Triton X-100), and wash buffer 5 (10 mM Tris-HCl). DNA libraries were prepared through the ChIPmentation method ^62^. Briefly, beads were resuspended in 30 μl of the tagmentation reaction buffer (10 mM Tris-HCl pH 8.0 and 5 mM MgCl_2_) containing 1 μl Tagment DNA Enzyme from the Nextera DNA Sample Prep Kit (Illumina) and incubated at 37°C for 10 min in a thermal cycler. The beads were washed twice with 150 μl cold wash buffer 1, incubated with elution buffer (10 mM Tris-HCl pH 8.0, 1 mM EDTA, 250 mM NaCl, 0.3% SDS, 0. 1 μg/μl Proteinase K) at 42°C for 30 min, and then incubated at 65°C for another 5 h to reverse cross-linking. DNA was purified with the MinElute Reaction Cleanup Kit (Qiagen) and amplified with NEBNext High-Fidelity 2× PCR Master Mix (NEB). Amplified DNA was purified by Agencourt AMPure XP (Beckman Coulter). Afterwards, DNA fragments in the 250- to 500-bp size range were prepared by agarose gel size selection. DNA libraries were adjusted to 5 nM in 10 mM Tris-HCl pH 8.0 and sequenced with an Illumina HiSeq 2500.

### Code availability: ChIP-seq and RNA-seq data

RNA-seq data from THY1^+^ spermatogonia, PS, and RS were downloaded from the Gene Expression Omnibus (accession number: GSE55060)^3^. ChIP-seq data for A-MYB and RNA-seq data from A-MYB mutant and control testes were downloaded from the Gene Expression Omnibus (accession number: GSE44690)^21^. ChIP-seq data for H3K4me3, and H3K4me2, and RNA-seq data from KIT^+^ spermatogonia, were downloaded from Gene Expression Omnibus (accession number: GSE89502)^40^. While generated for and analyzed in this study, our H3K27ac ChIP-seq data for wild-type PS and RS were initially introduced in another study that analyzed active enhancers on the sex chromosomes^23^; ChIP-seq data for H3K27ac from PS and RS, were downloaded from Gene Expression Omnibus (accession number: GSE107398)^23^. ChIP-seq data for H3K27ac from embryonic stem cells was downloaded from Gene Expression Omnibus (accession number: GSE29184)^63^. ChIP-seq data for H3K27ac from sperm were downloaded from Gene Expression Omnibus (accession number: GSE79230)^64^.

### ChIP-seq and RNA-seq data analysis

Data analysis for ChIP-seq was performed in the Wardrobe Experiment Management System (https://code.google.com/p/genome-tools/)^65^. Briefly, reads were aligned to the mouse genome (mm10) with Bowtie (version 1.2.0)^66^, assigned to RefSeq genes (or isoforms) using the Wardrobe algorithm, and displayed on a local mirror of the UCSC genome browser as coverage. ChIP-seq peaks for H3K27ac, H3K4me2, H3K4me3, and A-MYB were identified using MACS2 (version 2.1.1.20160309)^24^. Pearson correlations for the genome-wide enrichment of H3K27ac peaks among ChIP-seq library replicates were analyzed using SeqMonk (Babraham Institute). MAnorm, software designed for quantitative comparisons of ChIP-seq datasets^25^, was used to compare the genome-wide ChIP-seq peaks among stages in spermatogenesis. Unique peaks were defined using the following criteria: (1) defined as “unique” by the MAnorm algorithm; (2) *P*-value <0.01; (3) raw counts of unique reads >10. Common peaks between two stages were defined using the following criteria: (1) defined as “common” by MAnorm algorithm; (2) raw read counts of both stages >10. Average tag density profiles were calculated around transcription start sites for gene sets of somatic/progenitor genes, late spermatogenesis genes, constitutive active genes, and constitutive inactive genes as described previously^4^. Resulting graphs were smoothed in 200-bp windows. Enrichment levels for ChIP-seq experiments were calculated for 4-kb windows, promoter regions of genes (±2 kb surrounding TSSs) and enhancer regions. To normalize tag value, read counts were multiplied by 1,000,000 and then divided by the total number of reads in each nucleotide position. The total amount of tag values in promoter or enhancer regions were calculated as enrichment. The k-means clustering of differential enhancer peaks were analyzed using Cluster 3.0 software. The results were further analyzed using JavaTreeview software^67^ to visualize as heatmaps. MEME-ChIP ^68^ was used for motif discovery as described in the text. For all motif analyses, we used only peak regions (±250 bp from the peak summit) outside of ±1 kb from TSSs; we chose a maximum of 3,000 peak regions from the lowest *P*-values (*P* <0.01) via MAnorm analysis, and we extracted those sequences using the Table Browser^69^. The HOMER software package^36^ was used for motif enrichment analyses using a customized version of HOMER that employs a log base 2 scoring system and motifs contained in the Cis-BP motif database^37^. To identify SEs, H3K27ac ChIP-seq data were used with the same criteria and software as previously described^17,26^.

RELI (Regulatory Element Locus Intersection) analysis was performed as described previously^45^ In brief, genomic regions of interest (e.g., ChIP-seq peaks) were systematically aligned with a large collection of publicly available ChIP-seq data for various TFs in various cellular contexts largely taken from mouseENCODE, and the significance of the intersection of each dataset was calculated using RELI.

Analyses of RNA-seq data were performed in the Wardrobe Experiment Management System^65^. Briefly, reads were aligned by STAR (version STAR_2.5.3a)^70^ with “--outFilterMultimapNmax 1--outFilterMismatchNmax 2”. RefSeq annotation from the mm10 UCSC genome browser ^71^ was used. The --outFilterMultimapNmax parameter was used to allow unique alignments only, and the--outFilterMismatchNmax parameter was used to allow a maximum of 2 errors. All reads from the resulting .bam files were split for related isoforms with respect to RefSeq annotation. Then, the EM algorithm was used to estimate the number of reads for each isoform.

### Accession Codes

H3K27ac ChIP-seq data reported in this study are deposited to the Gene Expression Omnibus (GEO) under the accession number GSE130652. The following secure token has been created to allow review of record GSE130652 while it remains in private status: sbslqciwldgxpof

## Acknowledgements

We thank members of the Namekawa laboratory for discussion and helpful comments regarding the manuscript. Funding sources: The research project grant by the Azabu University Research Services Division, Ministry of Education, Culture, Sports, Science and Technology (MEXT)-Supported Program for the Private University Research Branding Project, (2016-2019), Grant-in-Aid for Research Activity Start-up (19K21196), and The Uehara Memorial Foundation Research incentive grant (2018) to S.M.; Lalor Foundation Postdoctoral Fellowship to A.S.; Albert J. Ryan Fellowship to K.G.A.; CCHMC Endowed Scholar and CpG grant awards to M.T.W.; National Institute of Health (NIH) DP2 GM119134 to A.B.; NIH R01 GM122776 and GM098605 to S.H.N.

## Author contributions

The manuscript was written by S.M., K.G.A., and S.H.N., with critical feedback from all other authors, and S.M. and S.H.N. designed the study. S.M. performed the H3K27ac ChIP-seq experiments. S.M. M.Y., X.C., A.S., K.G.A., M.T.W., A.B., and S.H.N. designed and interpreted the computational analyses; S.M. performed the majority of computational analyses. S.H.N. supervised the project.

