## Supplemental Figures for "Super-enhancer switching drives a burst in germline gene expression at the mitosis-to-meiosis transition"

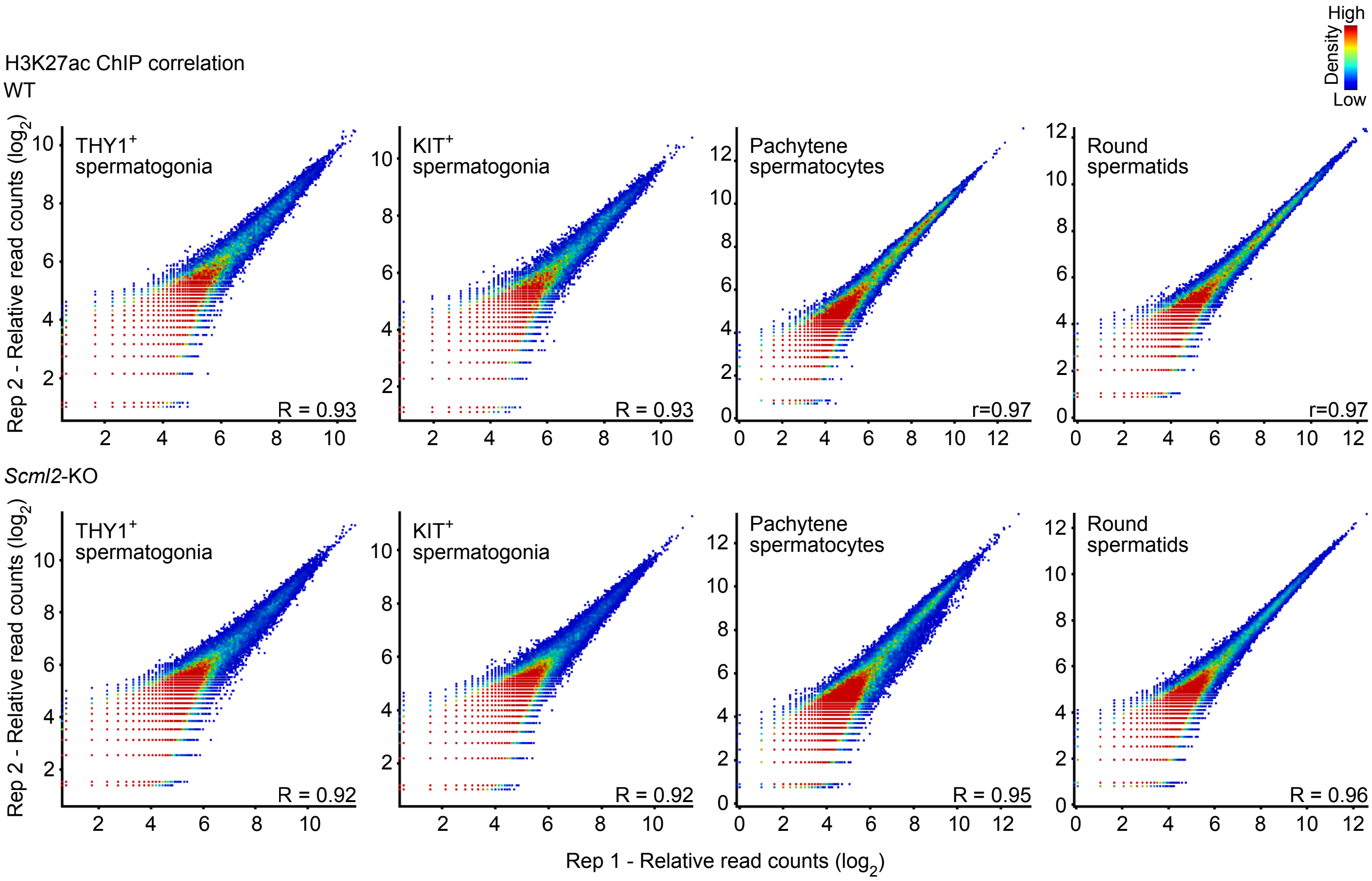


**Supplementary Figure 1. Biological replicates of H3K27ac ChIP-seq data.**
Two-dimensional scatter plots showing the reproducibility of H3K27ac ChIP-seq reads at individual peaks between biological replicates. Each peak was identified using MACS ($P < 1\times10\text{-}5$). Enrichment levels for H3K27ac ChIP-seq reads are shown in $Log_2$ RPKM values. The color scale indicates the density of H3K27ac ChIP-seq peaks. Pearson correlation values are shown. While generated for and analyzed in this study, our H3K27ac ChIP-seq data for wild-type PS and RS was initially introduced in another study that analyzed active enhancers on the sex chromosomes (these data are adapted from Adams et al. *PloS Genet* 2018).

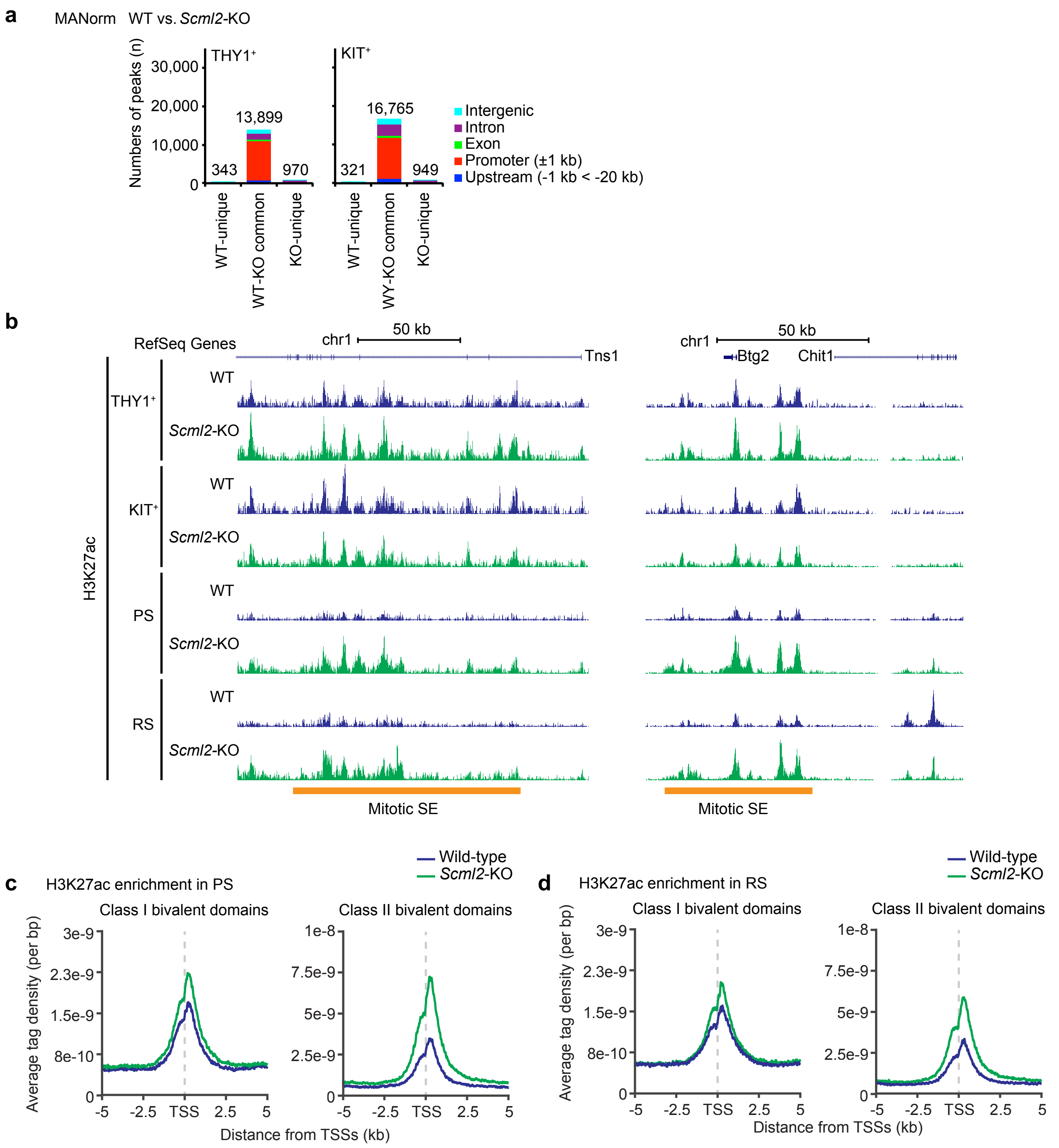


**Supplementary Figure 2. Comparison of H3K27ac ChIP-seq data between wild-type and *Scml2*-KO cells.**

(**a**) MAnorm analysis of H3K27ac peaks in THY1$^{+}$ and KIT$^{+}$ spermatogonia between wild-type and *Scml2*-KO. (**b**) Track view of H3K27ac ChIP-seq data on representative mitotic SEs in spermatogenesis. (**c**) Average tag density of H3K27ac ChIP-seq reads at genomic bivalent domains in PS and in RS.

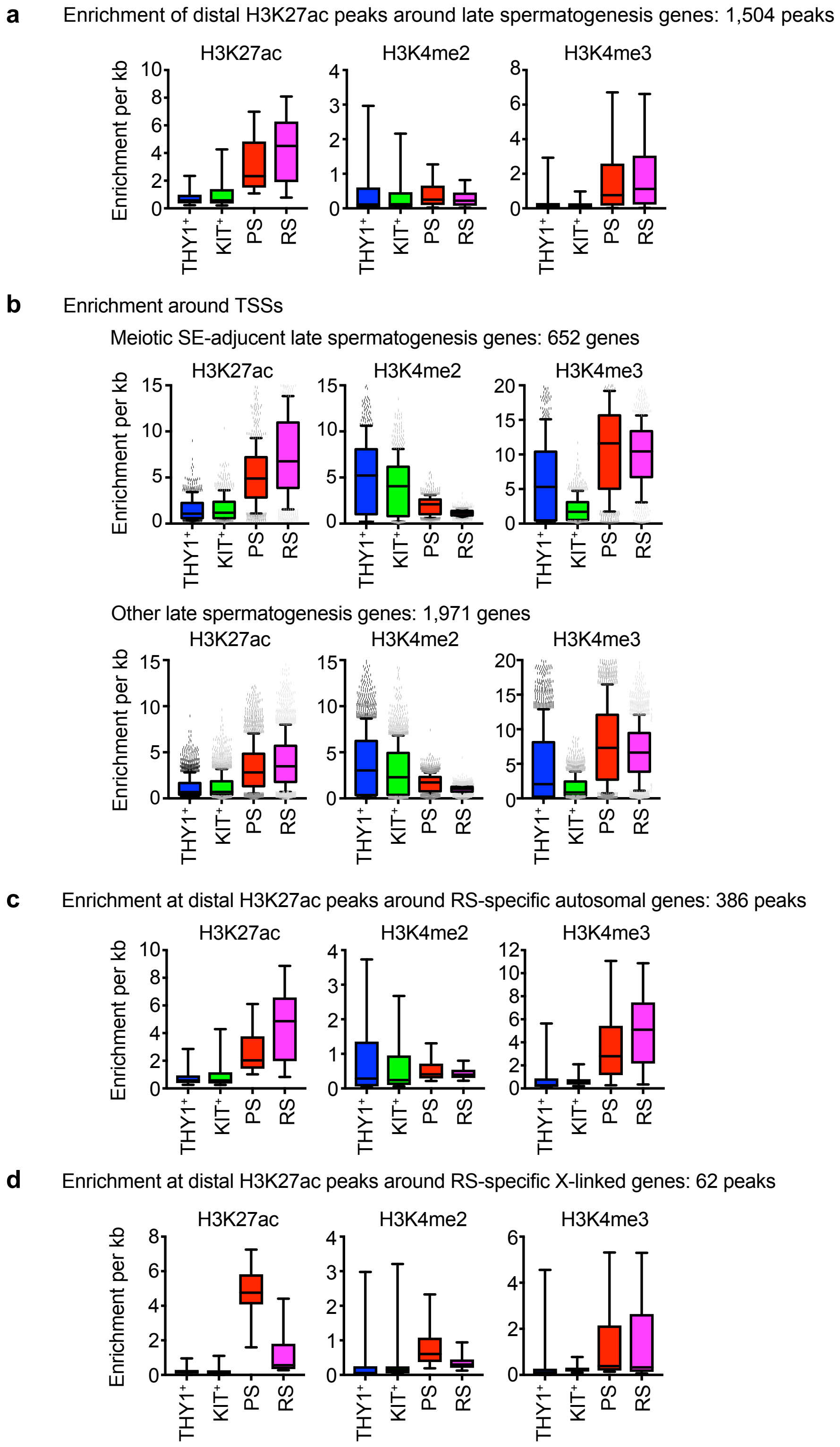


**Supplementary Figure 3. ChIP-seq read enrichment on various genomic loci**

(a-d) Box-and-whisker plots showing distribution of enrichment for ChIP-seq data for various genomic loci indicated. Central bars represent medians, the boxes encompass 50% of the data points, and the whiskers indicate 90% of the data points.
